# Feline mammary carcinomas display evidence of stemness, epithelial-mesenchymal plasticity, and metastasis-associated M2-like macrophage infiltration

**DOI:** 10.64898/2026.08.03.742512

**Authors:** Kimaya M. Bakhle, Sophie R. Nelissen, Gerald E. Duhamel, Anushka Dongre

## Abstract

Triple-negative breast cancer (TNBC) is the most aggressive subtype of breast cancer-a consequence of high proliferation rates, stemness, epithelial-mesenchymal plasticity, and immune evasion. Feline mammary carcinomas (FMCs) are spontaneous mammary gland cancers associated with high rates of metastasis and death. The aggressive biological behavior of FMCs, as well as their lack of expression of hormone receptors, make FMCs an excellent naturally-occurring model of human triple-negative breast cancer (TNBC). Therefore, we investigated whether FMCs of increasing histologic grade (I-III) harbor characteristics comparable to human TNBC. We performed immunohistochemistry using markers of proliferation (Ki67), stemness (Sox2), and epithelial-mesenchymal plasticity (E-cadherin, vimentin). We found that FMCs expressed high levels of Sox2 and vimentin together with a grade-associated increase in Ki67 expression. Apart from these cancer cell-intrinsic properties, knowledge on the tumor microenvironment of FMCs is limited. To this end, we assessed the presence of T-cells (CD3), total macrophages (Iba1), and immunosuppressive M2-like macrophages (CD204) in FMCs. High-grade FMCs showed increased M2-like macrophage infiltration. Moreover, samples with evidence of vascular invasion and lymph node metastasis displayed increased total and M2-like macrophage infiltration. These findings suggest that the presence of intratumoral macrophages is associated with the biological aggressiveness and metastatic potential of FMCs. Taken together, our results support FMCs as a translational model of human TNBC.

## Introduction

Triple-negative breast cancer (TNBC) is a subtype of breast cancer lacking the hormone receptors for estrogen (ER) and progesterone (PR), as well as human epidermal growth factor receptor 2 (HER2). Therefore, TNBC is unresponsive to endocrine therapies, such as tamoxifen, as well as HER2-targeted treatments, such as trastuzumab^1–4^. Additionally, TNBC mounts resistance to multiple forms of chemotherapies through upregulation of drug efflux pumps, acquisition of a dormant, stem-like phenotype, intratumoral heterogeneity, and metabolic changes^5–7^. TNBC also possesses high rates of disease recurrence and metastases to the brain and viscera^2,4,8,9^.

Feline mammary carcinomas (FMCs) are the third most common cancer in cats and display aggressive behavior and metastatic potential, leading to high morbidity and mortality rates^10^. Recently, FMCs have emerged as a naturally-occurring model of human TNBC based on a range of similarities. In addition to sharing common environmental risk factors and relative age of disease onset, FMCs have transcriptional signatures and genetic heterogeneity like human TNBC, including conserved *PIK3CA* and *TP53* driver mutations^10,18^. The majority of FMCs lack expression of ER, PR, and HER2, making them an ideal model for human TNBC^11–14^. Moreover, histologic grading of FMCs follows a modified version of the Elston and Ellis grading system used for human breast cancer. Finally, histologic grade is a strong prognostic indicator of survival in both species^10,16,17^. To understand whether FMCs conserve other cancer cell-intrinsic and -extrinsic features of human TNBC, we used immunohistochemistry to assess patient-derived FMC samples with a panel of cellular markers. Cancer cell-intrinsic features of interest included proliferation and stemness, as well as EMP. TNBC possesses high proliferation rates, as measured by nuclear Ki67 immunohistochemistry^19,20^. TNBC also displays features of cancer stem cells (CSCs), which have unique self-renewal and tumor-initiating capacity. Specifically, TNBC cells share transcriptional signatures with CSCs as well as increased ability to form tumors in mice when implanted at low numbers in laboratory mice^21–25^. CSCs express several transcription factors, including Sox2, which maintain a stem-like state and capacity for self-renewal^26,27^. CSCs also bear similarities to dormant/persistent cancer cells, which drive disease recurrence after treatment^23^. Finally, CSCs contribute to intratumoral heterogeneity through emergence of resistant cancer cell subpopulations^23,28,29^.

One mechanism of CSC generation in TNBC is EMP activation^25,30,31^. EMP is an epigenetic program activated in carcinoma cells to promote migratory and invasive potential and resistance to multiple therapeutic modalities^31,32^. This process enables epithelial cells, which express markers such as E-cadherin, to transition to a series of hybrid or quasi-mesenchymal phenotypic states, marked by upregulation of mesenchymal markers including vimentin. EMP is driven by families of transcription factors including Snail, Slug, Zeb, and Twist^33–39^. EMP transcription factor upregulation has been previously demonstrated in a small batch of FMC samples^15^. EMP activation leads to acquisition of a spindle-shaped morphology, loss of cell-cell adhesion proteins, and upregulation of matrix metalloproteases (MMP), promoting an invasive phenotype^35,40–42^. This ultimately initiates the invasion-metastasis cascade, whereby carcinoma cells detach from the primary tumor and travel through blood and lymphatic circulation to colonize distant metastatic sites^33,42–45^. In addition to these cancer-cell intrinsic consequences, EMP also modulates the tumor microenvironment (TME) to promote immunosuppression and immunotherapy resistance^33,46^.

TNBC has been shown to assemble an immunosuppressive TME, in part due to EMP activation and hypoxic conditions^47–52^. The TME is a diverse and dynamic collection of cancer, immune, and stromal cells. Intratumoral immune cells include innate cells, such as macrophages, natural killer (NK) cells, neutrophils, dendritic cells, and monocytes, and adaptive cells including T-cells and B-cells. Presence of intratumoral T-cells is associated with positive prognosis and improved response to immunotherapy^53–60^. Immune cells within the TME interact with cancer cells through paracrine and juxtacrine signaling to promote or restrain tumorigenesis. For example, CD8^+^ T-cells, CD4^+^ Th1 cells, B-cells, NK cells, and M1-like macrophages recognize and kill cancer cells. Conversely, CD4^+^ regulatory T cells and M2-like macrophages promote tumor growth by inhibiting anti-tumor immune cells and releasing paracrine factors that act directly on cancer cells^53,55–58,61–63^.

To determine whether FMCs share cancer cell-intrinsic features of TNBC, we assessed formalin-fixed, paraffin-embedded (FFPE) FMC samples submitted for routine histopathology evaluation for the presence of markers of proliferation (Ki67), stemness (Sox2), and EMP (E-cadherin, vimentin). We show that FMCs expressed high levels of Sox2 and exhibit heterogeneous expression of vimentin, suggesting EMP activation and stem-like properties. Additionally, Ki67 index was positively associated with grade, consistent with higher proliferative activity of high-grade FMCs. We then investigated the TME of FMCs and found few tumor-infiltrating T-cells, which may reflect an immune-excluded TME. The degree of macrophage infiltration was associated with increased grade and evidence of metastasis, suggesting that intratumoral macrophages are associated with FMC invasion and metastasis. Taken together, these findings support FMCs as a naturally-occurring model of human TNBC and identify a potential role for tumor-associated macrophages in driving disease progression.

## Results

### FMCs show high levels of stemness marker Sox2 and a grade-associated increase in Ki67 proliferative index

We first investigated cancer cell-intrinsic properties of FMCs including proliferation and stemness and their association with histologic grade. A total of 32 archived FMCs of histologic grade I to III were selected from a database of client-owned female cats. The average age of the cats 9.75 years and all but four cats were spayed **(Table S1).** Information regarding birth control administration in unspayed cats was not available. There was no difference in age between histologic grades **(Fig S1A)**.

We first calculated the Ki67 index for all samples by dividing the number of Ki67^+^ tumor cells by the number of total tumor cells. We found that Ki67 proliferative index was significantly increased in grade III tumors compared to grade I tumors **(Fig 1A-B)**. This was in alignment with the mitotic index, which is calculated by manual counting of mitotic figures within a 2.37 mm^2^ field of corresponding H&E-stained tissue sections and showed a trend towards increased expression in grade III FMCs **(Fig 1B).** Moreover, Ki67 index and mitotic index were positively correlated **(Fig 1C).** Finally, all samples showed high levels of Sox2 expression in cancer cells, indicating that FMC cells have stem-like properties **(Fig 1A, 1D)**.

**Figure 1.**
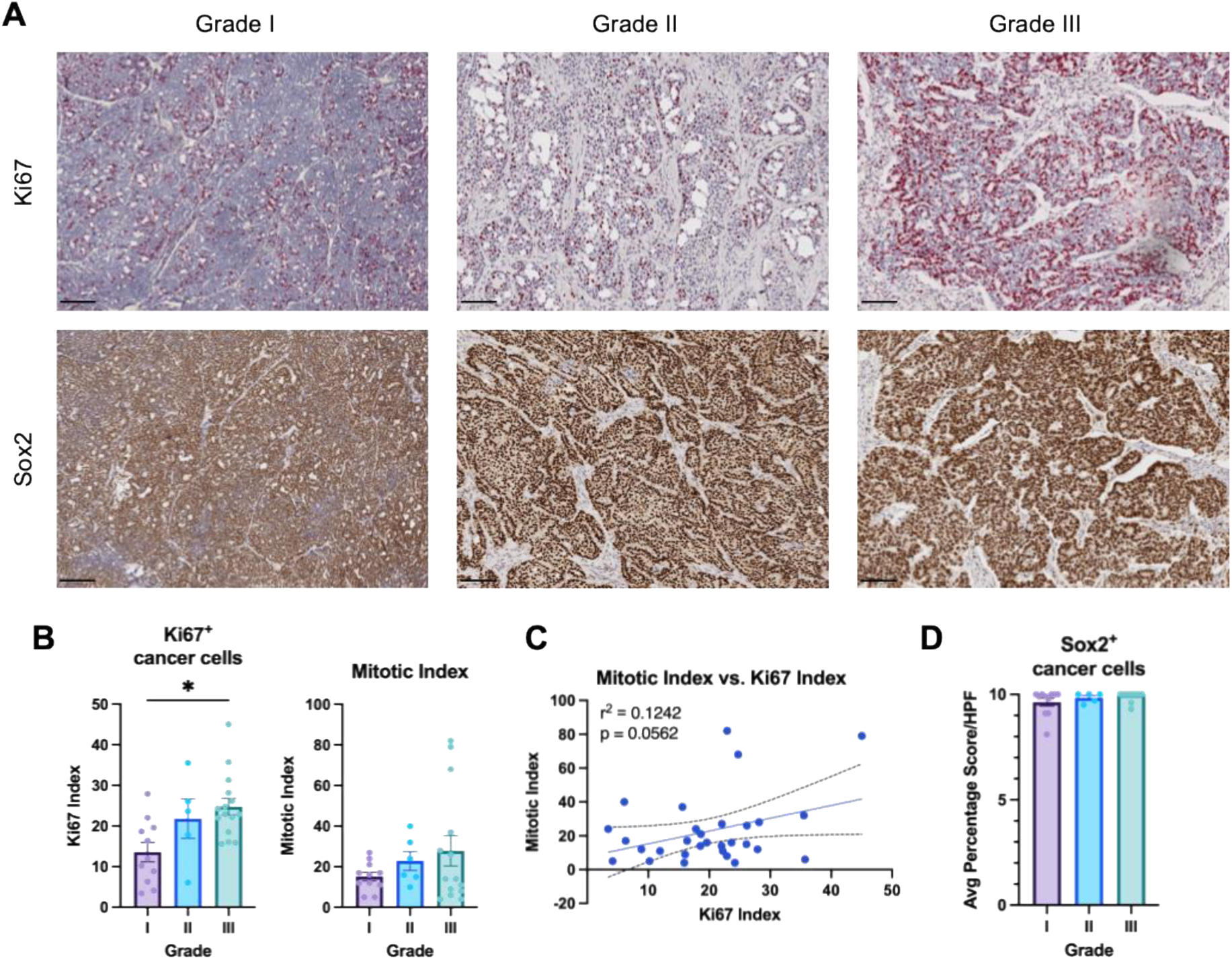
Ki67 and Sox2 labeling in feline mammary carcinomas (FMCs). **(A)** Representative images of archived formalin-fixed paraffin-embedded (FFPE) FMC specimens acquired from the New York State Veterinary Diagnostic Laboratory and IHC labeled with the proliferation marker Ki67 and the stemness marker Sox2. **(B)** Quantification of cancer cell-specific Ki67 labeling and mitotic index in FMCs. **(C)** Spearman correlation analysis of Ki67 and mitotic indices for FMC samples. **(D)** Quantification of cancer cell-specific Ki67 and Sox2 labeling in FMCs. All graphs are plotted as the mean value with error bars indicating the standard error of the mean. Data analyzed using nonparametric one-way ANOVA: * p < 0.05, ns (p ≥ 0.05) for remaining comparisons. Images taken at 10x magnification, scale bars indicate 100 µm.

### FMCs show evidence of epithelial-mesenchymal plasticity independent of grade

Next, we wanted to determine whether EMP is associated with increased grade in FMCs, as we previously documented in canine mammary carcinomas^64,65^. We found that FMCs of all grades showed high E-cadherin membranous expression **(Fig 2A-B)**. There was a trend towards a decrease in the composite E-cadherin score in grade III tumors, suggesting that the intensity of E-cadherin expression may be decreased in high-grade tumors **(Fig 2B)**. The expression of vimentin was not significantly different between tumors of different grades. Moreover, vimentin expression showed heterogeneity between samples and within an individual sample **(Fig 2A, 2C)**. E-cadherin and vimentin co-expression was observed not only in spindle-shaped cancer cells as canonically expected **(Fig 2D)**, but also in cuboidal cells **(Fig 2E)**. This suggests that FMCs contain a spectrum of EMP states, including partial or hybrid phenotypes, that vary in cellular morphology, independent of histologic grade.

**Figure 2.**
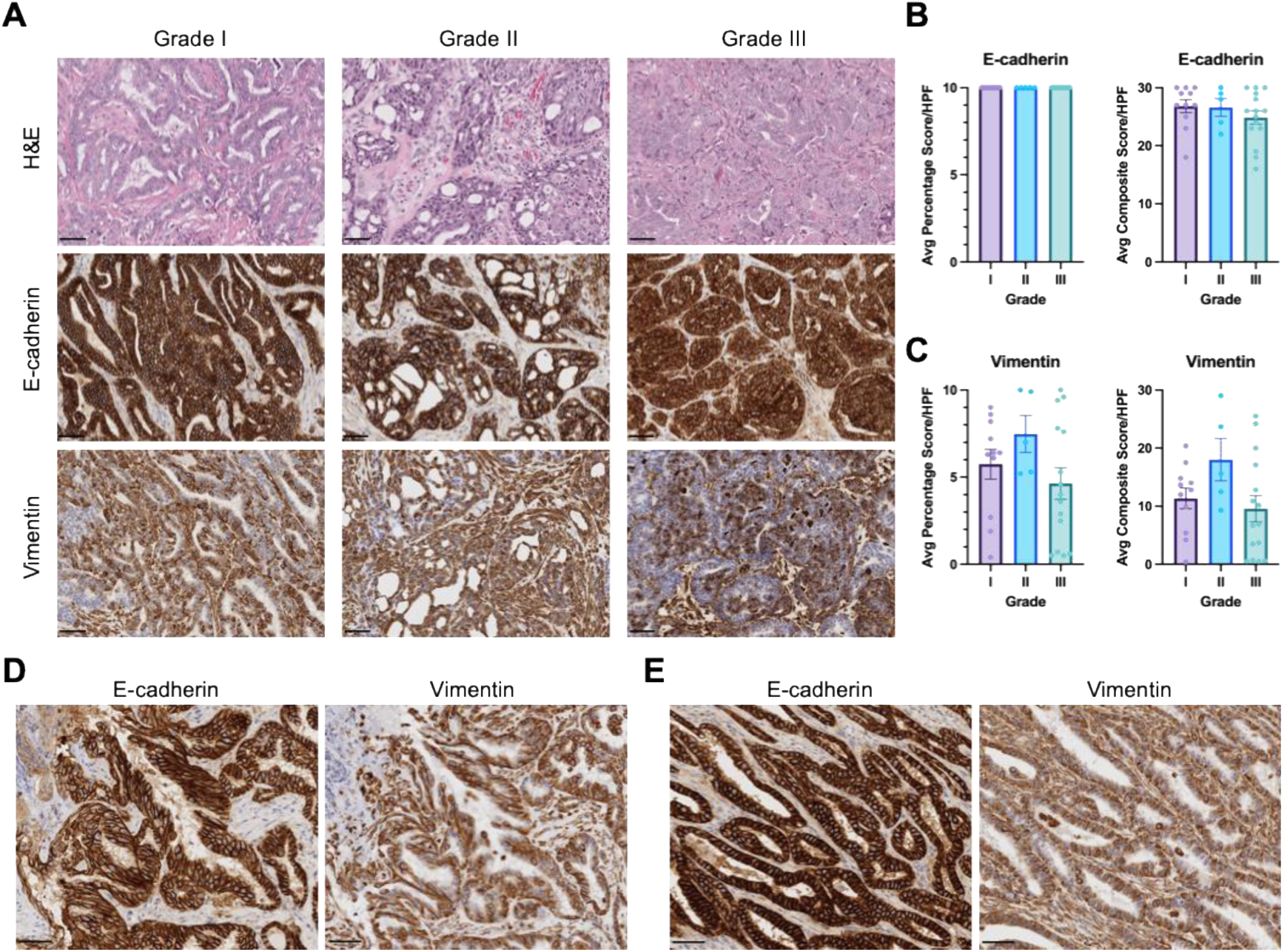
Expression of epithelial-mesenchymal plasticity (EMP) markers in FMCs. **(A)** Representative images of FMC specimens IHC labeled with the epithelial marker E-cadherin and the mesenchymal marker vimentin. **(B-C)** Semi-quantitative analysis of cancer cell-specific E-cadherin and vimentin expression, as percentage of cells and a composite score accounting for both percentage and intensity of labeling across 10 high-power fields (HPF, 2.37mm²). All graphs are plotted as the mean value with error bars indicating the standard error of the mean. Data analyzed using nonparametric one-way ANOVA: ns (p ≥ 0.05) if not indicated. Images taken at 20x magnification, scale bar indicates 50 µm. ROI = region of interest. **(D)** Representative images of E-cadherin and vimentin co-expression in spindle-shaped cancer cells. **(E)** Representative images of E-cadherin and vimentin co-expression in cuboidal cancer cells. Scale bars indicate 50 µm.

### High-grade FMCs preferentially recruit CD204^+^ M2-like macrophages

In addition to the cancer cell-intrinsic properties of proliferation, stemness, and EMP activation, we sought to profile immune cells within the TME of FMCs. We first assessed expression of the T-cell marker CD3 and found that there were few tumor-infiltrating T-cells in FMCs of all grades **(Fig 3A-B)**. We also observed phenotypes of T-cell-infiltrated and -excluded tumors like those described in murine and human tumors **(Fig 3C)**^66–71^.

**Figure 3.**
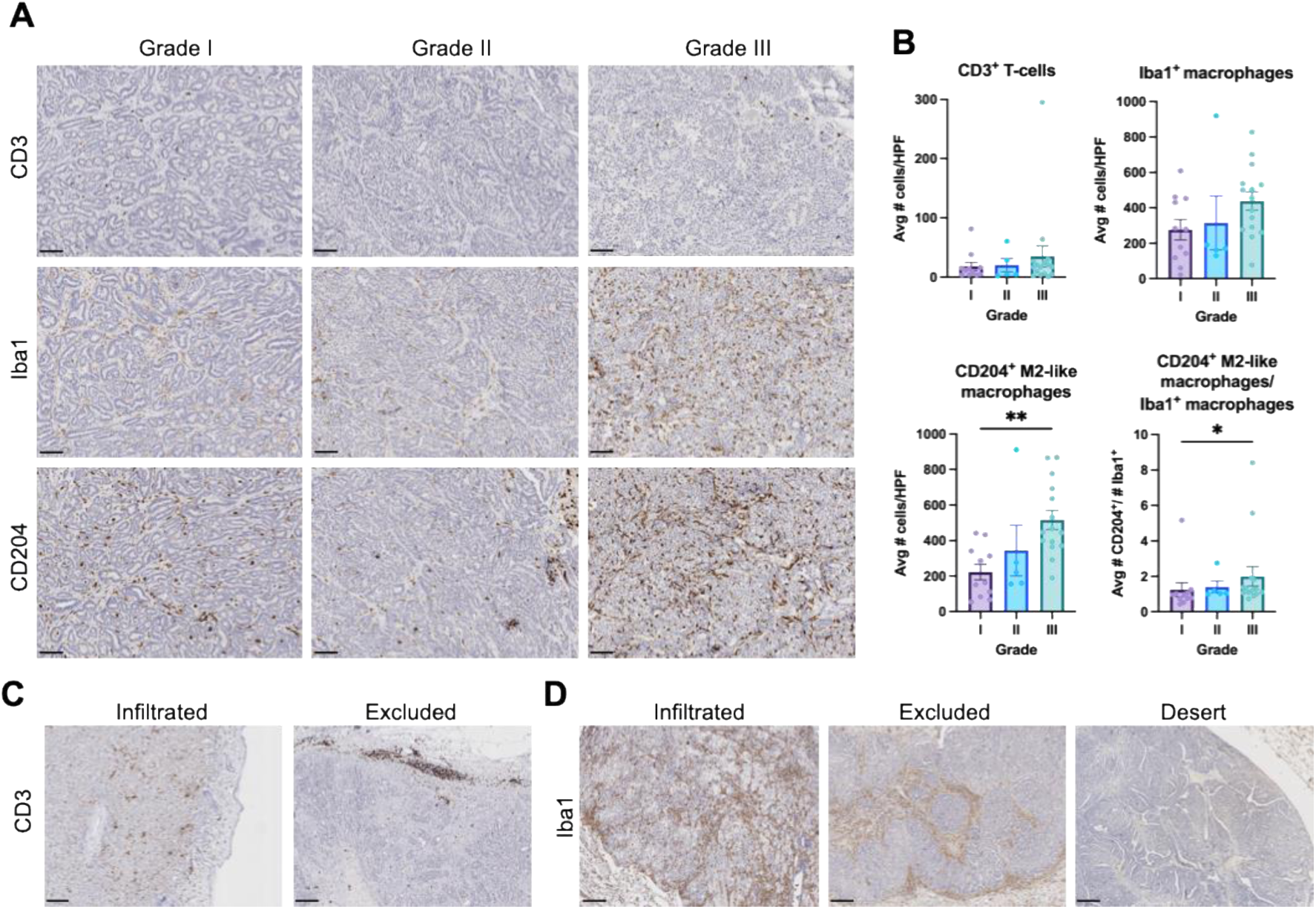
T-cell and macrophage infiltration in FMCs. **(A)** Representative images of FMC specimens IHC labeled with the T-cell marker CD3, the pan-macrophage marker Iba1, and the M2-like macrophage marker CD204. **(B)** Quantification of intratumoral T-cells and macrophages across 10 high-power fields (HPF). All graphs are plotted as the mean value with error bars indicating the standard error of the mean. Data analyzed using nonparametric one-way ANOVA: ** p < 0.01, * p < 0.05, ns (p ≥ 0.05) if not indicated. **(C)** Representative images of T-cell-infiltrated and -excluded FMCs. **(D)** Representative images of macrophage-infiltrated, -excluded, and desert FMCs.

We next quantified tumor-infiltrating macrophages using the pan-macrophage marker Iba1, as well as the M2-like macrophage marker CD204. We found that there was a trend towards increased total macrophages in grade III tumors compared to those of grade I **(Fig 3A-B).** Both the number of CD204^+^ M2-like macrophages as well as the ratio of CD204^+^ M2-like macrophages/Iba1^+^ total macrophages was significantly increased in grade III tumors **(Fig 3A-B)**. Like the patterns of T-cell infiltration, we observed phenotypes of macrophage-infiltrated, -excluded, and desert tumors **(Fig 3D)**. Together, these findings suggest that M2-like macrophage infiltration is associated with increased histologic grade and may be a driver of more aggressive tumor biological behavior.

### Vascular invasion and lymph node metastasis are associated with increased macrophage infiltration in the primary tumor

Since lymphovascular invasion is a component of histologic grading in FMCs, we next asked whether this was associated with macrophage infiltration. While none of the grade I or grade II tumors had evidence of vascular invasion, 11 out of 14 grade III cases had vascular invasion **(Fig S1B).** We found that the number of both Iba1^+^ macrophages and CD204^+^ M2-like macrophages were significantly increased in tumors with evidence of vascular invasion **(Fig 4A-B).**

**Figure 4.**
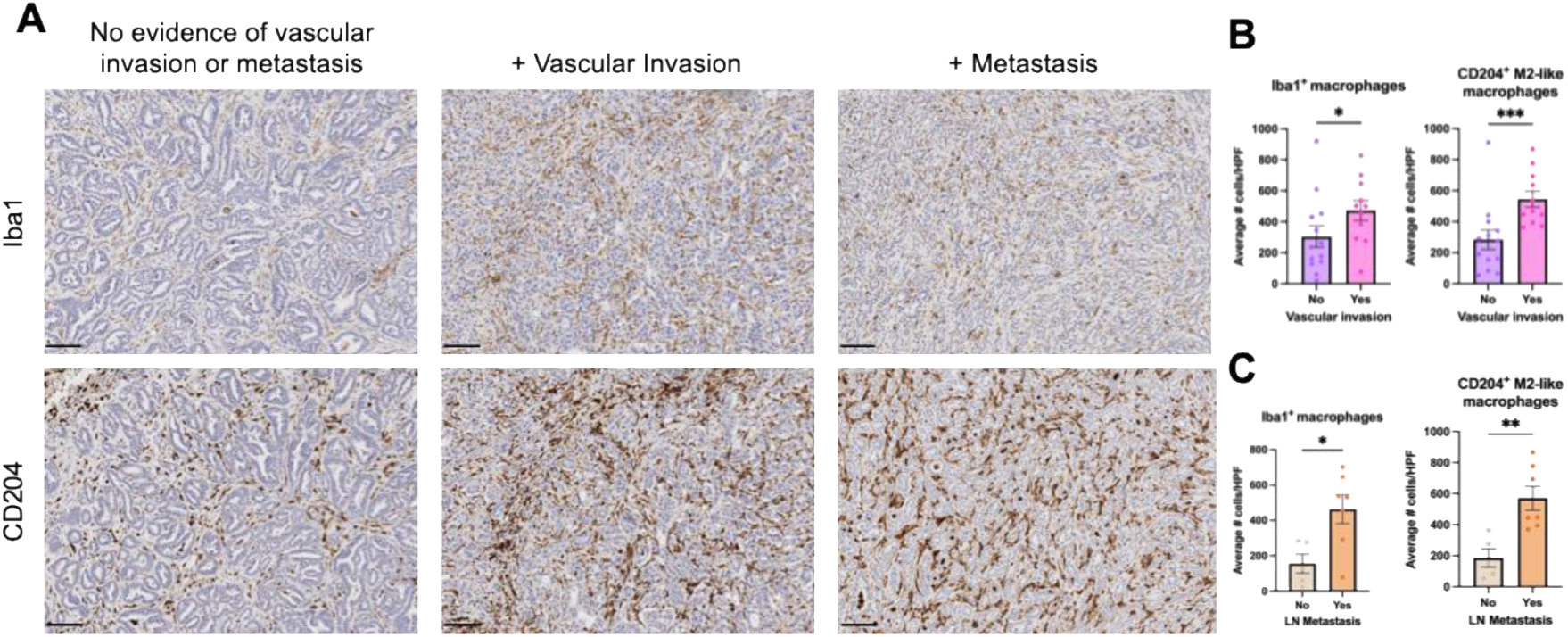
FMCs with evidence of vascular invasion and metastasis display increased macrophage infiltration. **(A)** Representative images of Iba1 and CD204 IHC labeled FMCs with or without evidence of vascular invasion and metastasis. Quantification of intratumoral macrophages across 10 high-power fields (HPF) based on presence of vascular invasion **(B)** or lymph node metastasis **(C)**. All graphs are plotted as the mean value with error bars indicating the standard error of the mean. Data analyzed using nonparametric one-way ANOVA: *** p < 0.001, ** p < 0.01, * p < 0.05.

Finally, because lymphovascular invasion precedes metastatic dissemination, we wanted to determine if cases with evidence of metastasis also had increased macrophage infiltration. Fewer grade I and grade II cases had lymph nodes (LN) available for metastatic evaluation, but none of these cases had evidence of metastasis. By contrast, seven out of the eight LN evaluated from grade III tumors contained neoplastic cells **(Fig S1C).** Indeed, these seven primary tumor samples showed significantly increased infiltration of Iba1^+^ macrophages and CD204^+^ M2-like macrophages **(Fig 4A, 4C).** In FMCs without evidence of lymphovascular invasion or LN metastasis, the intratumoral macrophages were restricted to the stroma surrounding clusters of neoplastic cells. Conversely, in FMCs with lymphovascular invasion or LN metastasis, macrophages were in direct contact with cancer cells. These results provide supporting evidence for a role of intratumoral macrophages in driving invasive and metastatic properties of FMCs.

## Discussion

In this study, we identified several conserved features between human TNBC and FMCs including high proliferative rates, stemness, and EMP. The association between increased Ki67 index and grade is in alignment with the “mitotic count” criterion already used to assign histologic grade to FMCs (≤ 62 or > 62 mitotic figures in 10 consecutive 40x fields)^17^. Ki67 labeling in FMCs has been performed by other groups, but direct Ki67 proliferative index association with histologic grade has not been previously described^72–75^. Though calculating a mitotic index was previously considered tedious, we here demonstrate that combining the use of a pixel classifier and cell detection tool is sufficient to adequately calculate a Ki67 index in the open-source software QuPath^76^. Based on these findings, Ki67 index of FMCs may realistically serve as a useful prognostic tool for FMCs.

Additionally, we assessed FMC samples for Sox2 expression to assess cancer cell stemness in FMCs. We found that all tumors expressed nuclear Sox2 with similar intensity. Some grade I tumors showed Sox2 expression in <100% of cancer cells, but the majority of tumors showed ubiquitous and consistent cancer cell-intrinsic Sox2 expression throughout the tumor. Previous reports have identified Sox2 expression in less than 50% of FMC samples^77^, which suggests that our cohort may represent a subset of especially stem-like FMCs. Conversely, this discrepancy may also reflect geographic (USA vs. France), environmental, and “breed”-specific variations (American vs. European domestic longhair and domestic shorthair cats).

Since human TNBC is known to activate EMP^30^, we investigated the expression of the epithelial marker E-cadherin and mesenchymal marker vimentin in FMCs. We found that all samples maintained E-cadherin expression, but there was heterogeneous vimentin expression between different tumors and within an individual tumor. Expression of these markers was not associated with histologic grade, unlike our previous observation with canine mammary carcinomas^64,65^. This suggests that FMCs do activate EMP, but not in a uniform fashion across cancer cells. We have observed similar segregation of epithelial and quasi-mesenchymal cancer cells in murine models of breast cancer as well. In this case, minority fractions of quasi-mesenchymal cells were sufficient to drive immunosuppression in the entire heterogeneous tumor^47^. This suggested that EMP activation in a subset of cancer cells may promote progression of the entire FMC. Other groups have investigated E-cadherin and vimentin expression in FMCs separately. In this context, they have identified that hormone receptor-negative FMCs express vimentin^14,15^. Loss of E-cadherin has been associated with metastasis of FMCs^78,79^. We found that a E-cadherin expression was not significantly decreased in grade III FMCs, of which, a high percentage were metastatic. Previous studies investigating E-cadherin and vimentin expression in FMCs have not characterized the co-expression of these two markers. Using immunohistochemistry of serial tumor sections, we identified E-cadherin^+^ Vimentin^+^ neoplastic cells. These cells may represent a hybrid epithelial-mesenchymal population of cancer cells. This hybrid phenotype has been shown to be extremely tumorigenic and metastatic in murine models^37–39^, which suggests the hybrid state could be a factor driving the well-documented aggressive biological behavior of FMCs. Future studies should examine the expression of additional EMP markers, as well as EMP-inducing TFs in FMCs.

We next investigated the FMC TME and found that high-grade FMCs recruit increased numbers and proportions of CD204^+^ M2-like macrophages, similar to our previous findings in naturally-occurring canine mammary carcinomas^64^. Findings from De Haes et al. support increased macrophage recruitment in grade III FMCs, but in their study, the macrophages were restricted to the tumor periphery, whereas we found dense infiltration of macrophages throughout FMCs^80^. Since mitotic index is an important grading criterion of both canine and feline mammary carcinomas, it is possible that macrophage recruitment has a pro-proliferative effect in both species. We did not find associations between T-cell infiltration and grade; however, most FMCs had few tumor-infiltrating T-cells. Additionally, since we only labeled tumors for CD3, it is unknown whether these are immunosuppressive Tregs or immunostimulatory cytotoxic lymphocytes or Th1 cells, or if they express markers of dysfunction (e.g. PD-1, CTLA-4).

Finally, we investigated whether macrophage recruitment could be linked to another component of FMC grading: vascular invasion. Indeed, both evidence of vascular invasion and distant metastasis were both associated with significantly increased numbers of total and M2-like macrophages. Tumor-associated macrophages (TAMs) can secrete a variety of pro-tumorigenic and pro-angiogenic factors. For example, secretion of TGF-β and IL-6 by TAMs can induce EMP in surrounding cancer cells, thereby promoting invasion and metastasis^33^. Additionally, secretion of VEGFA by TAMs leads to tumor neovascularization, creating additional routes for metastasis^81,82^. Finally, TAMs produce matrix metalloproteases such as MMP9 and MMP19, which degrade the extracellular matrix (ECM) to promote cancer cell migration^82^. Therefore, future work should profile these TAMs and their ability to modulate the surrounding TME through EMP activation, angiogenesis, and ECM remodeling.

In summary, we have found several novel features of FMCs that may promote tumorigenesis in a manner similar to human TNBC. Characteristics of EMP and stemness in cancer cells may promote invasion, metastasis, disease recurrence, and immune evasion in FMCs. Additionally, the TAM-rich TME of high-grade FMCs likely promotes tumor progression and metastatic dissemination through paracrine effects on surrounding cancer, immune and stromal cells. These findings support studying FMCs as a spontaneous animal model of human TNBC and provide motivation to investigate TAM-driven invasion and metastasis in FMCs.

## Materials and Methods

### Sample selection and histologic analysis

Pathology archives of the New York State Animal Health Diagnostic Center (NYS-AHDC) at Cornell University were retrospectively reviewed from 2019 to 2025 to identify FMCs. In each case, tumor tissue samples were submitted as part of routine clinical care by referring veterinary practitioners of client-owned animals. All cases were initially evaluated by the attending American College of Veterinary Pathology (ACVP) board-certified veterinary pathologist at the time of submission. Per internal protocol, all samples received by the NYS-AHDC are the property of the NYS-AHDC and do not need owner consent for research use. The use of archived diagnostic material of client-owned animals for the purpose of this study is considered exempt from review by Cornell University’s Institutional Animal Care and Use Committee.

Thirty-two formalin-fixed paraffin-embedded (FFPE) feline mammary carcinoma (FMC) cases were identified for further analysis by immunohistochemistry **(Table S1)**. All available histopathologic slides were reviewed and graded from I to III, by a ACVP board-certified veterinary pathologist (G.E.D.) using the revised Elston and Ellis system for prognosis of FMC, originally developed for human breast cancer, and based on criteria defined by Mills et al.^17^, including mitotic count exceeding 62 per 2.37 mm^2^ field of HE-stained tissue sections (mitotic index), presence of lymphovascular invasion, and greater than 5% abnormal nuclear form. Of the 32 FMCs, 11 were classified as grade I, 6 as grade II, and 15 as grade III cases.

### Immunohistochemical labeling and quantification

For immunohistochemistry (IHC), formalin-fixed paraffin-embedded (FFPE) tissue sections were processed at the NYS-AHDC, Cornell University (Ithaca, NY, USA) using standard protocols and an automated IHC stainer (Leica Bond-Max, Leica Biosystems, Buffalo Grove, Illinois, USA). Briefly, serial 4-µm tissue sections mounted on charged slides were deparaffinized (AR9222, Bond dewax solution; Leica), and after heat epitope retrieval (AR9640, Bond epitope retrieval solution 2; Leica), the slides were incubated with the respective primary antibody followed by polymeric horseradish peroxidase (DS9800; Bond polymer refine detection; Leica) linker antibody conjugate detection system, and hematoxylin counterstain (DS9390; Leica) as previously described ^63^.

The following primary antibodies were used at the indicated concentrations: Ki67 (clone MIB-1, Dako M7240, RRID: AB_2142367, 1:50), Sox2 (clone 20G5, ThermoFisher MAI-014, RRID: AB_2536667, 1:500), E-cadherin (clone DECMA-1, Millipore MABT26, RRID: AB_10807576, 1:75), vimentin (clone SRL33, Leica PA0640-u, Ready to use), CD3 (clone LN10, Leica PA0553, RRID: AB_10554601, Ready to use), Iba1 (polyclonal, Wako 019-19741, RRID: AB_839504, 1:5000), CD204 (clone SRA-E5, Cosmo Bio KAL-KT022, 1:500). IHC staining was conducted as previously described^64,83–85^. Appropriate positive control tissues were included for each marker **(Fig S2A).** Additionally, tumor tissues labeled with secondary antibodies only were included to account for non-specific labeling **(Fig S2B)**. Labelled IHC and routine H&E-stained slides were scanned at 40x using an Aperio CS2 ScanScope (Visa, California, USA) provided by the Cornell University Department of Biomedical Sciences. IHC scoring and immune cell quantification was performed using QuPath software designed for digital pathology image analysis^76^.

To calculate the Ki67 index for each tumor, we trained a pixel classifier to differentiate tumor and stromal regions (QuPath). We then used the positive cell detection feature to quantify the number of Ki67^+^ tumor cells. The Ki67 index was calculated by dividing the number of Ki67^+^ tumor cells by the total number of tumor cells, as quantified by QuPath.

For E-cadherin, vimentin, and Sox2 IHC, 10 randomly selected 490 μm x 490 μm (2.37mm²) regions of interest (ROIs) were manually scored for % of cells expressing the marker (percentage score; 0-10 scale), staining intensity (0-3 scale), and notes about the localization of the marker and morphology of the stained cells were recorded. For E-cadherin and vimentin labeling, the two numeric scores (percentage score and staining intensity) were then multiplied to generate a composite score. The average percentage score and composite score for each FMC were plotted by grade.

For CD3, Iba1, and CD204, the number of positively labeled cells were quantified across 10 randomly selected 490μmx490μm (2.37mm²) ROIs using QuPath’s positive cell detection feature. The average number of positive cells across all 10 ROIs was plotted by grade for each tumor.

### Statistical Analysis

All statistical analyses were performed using the GraphPad Prism v10.4.1 software. All count data are plotted as mean with standard error of the mean (SEM) and analyzed using nonparametric ordinary one-way ANOVA. Pearson correlation analysis was used to correlate Ki67 and mitotic indices. Asterisks indicate statistical significance where * = p<0.05 and ** = p<0.01. Comparisons are not significant (p≥0.05) if not indicated. All data points were included for each experiment performed.

### Data Availability

All data generated and analyzed during this study are included in this article.

## Acknowledgments

We thank all staff in the Cornell University Animal Health Diagnostic Center for automated immunohistochemical labeling of FMC samples. We thank Marco Hiller for assistance with the Aperio ScanScope Unit used to digitally scan slides. Funding for this study was provided by the Cornell Feline Health Center.

