## Supplementary materials for "Feline mammary carcinomas display evidence of stemness, epithelial-mesenchymal plasticity, and metastasis-associated M2-like macrophage infiltration"

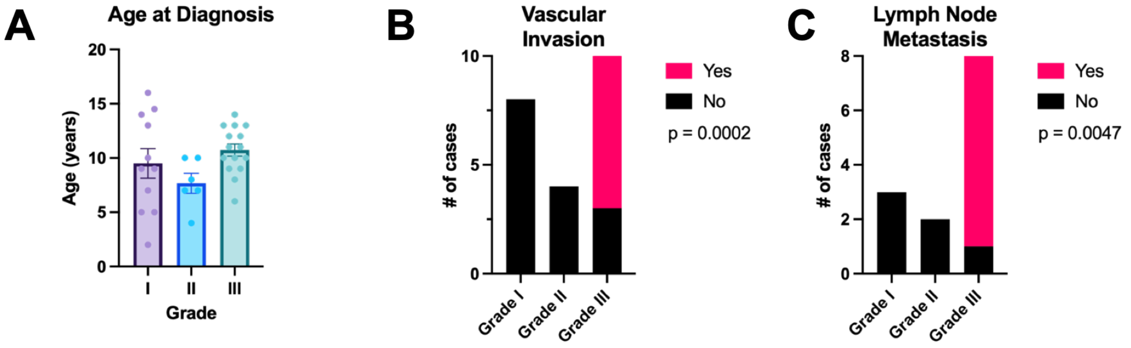


**Figure S1. Clinical parameters of FMC cases. (A)** Age at diagnosis of FMC cases, according to grade. Number of cases with evidence of vascular invasion **(B)** and lymph node metastasis **(C)**. Data analyzed using Fisher's exact test.

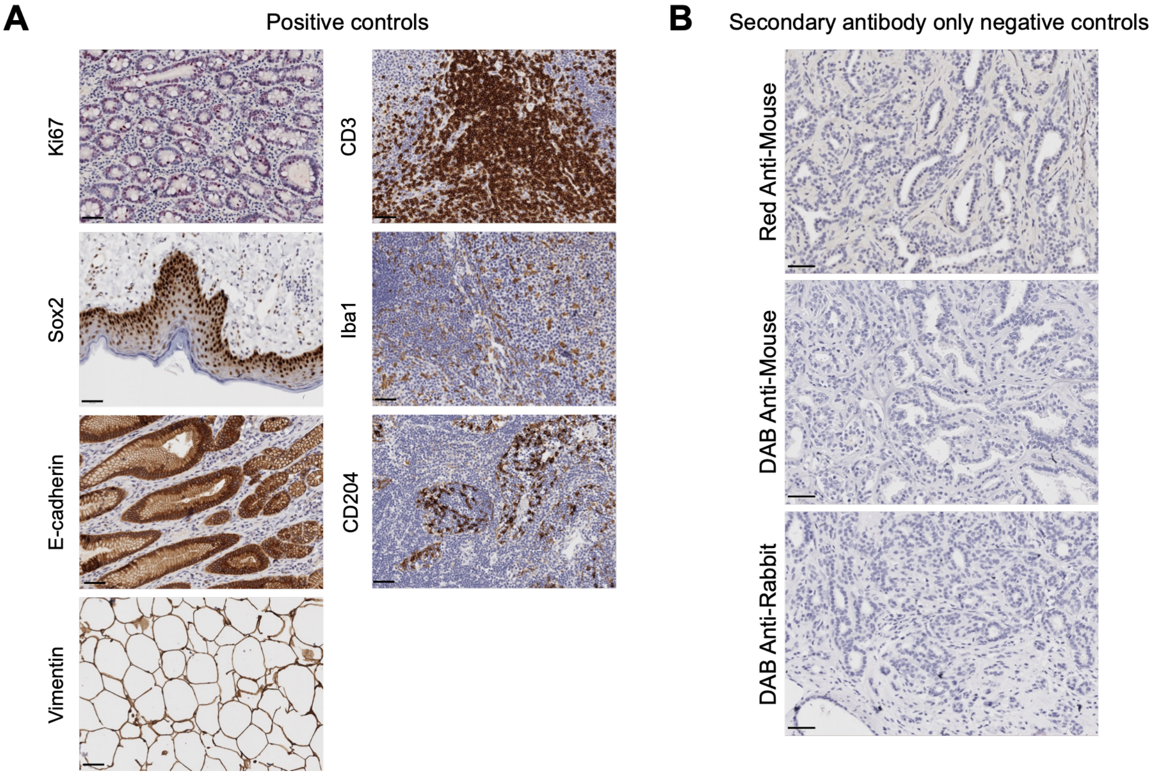


**Figure S2. Positive and negative controls for immunohistochemistry markers. (A)** Biological positive controls for all antibodies used in immunohistochemical labeling. Ki67: colon crypts, Sox2: skin, E-cadherin: gastric epithelium, vimentin: adipose tissue, CD3: lymph node, Iba1: lymph node, CD204: lymph node. **(B)** Secondary antibody-only negative controls for antibodies used immunohistochemical labeling.

**Table S1. Signalment and clinical parameters of FMC cases.**

| Sample | Sex | Age | Breed | Mammary carcinoma subtype | Grade | Mitotic Index | Vascular invasion? | LN evaluated? | Metastasis? |
|---|---|---|---|---|---|---|---|---|---|
| 1 | FS | 13 | DSH | Comedocarcinoma | I | 5 | No | No | |
| 2 | FS | 14.5 | DSH | Tubulopapillary carcinoma | I | 16 | No | No | |
| 3 | FS | 2 | DSH | Cribriform carcinoma | I | 5 | No | No | |
| 4 | FS | 9 to 13 | Siamese | 1. Simple carcinoma, 2. Complex carcinoma | I | 11 | No | Yes | No |
| 5 | F (Spayed but ovarian remnant identified) | 16 | DSH | Papillary carcinoma | I | 27 | No | No | |
| 6 | FS | 14 | Domestic | Tubulopapillary carcinoma | I | 24 | No | No | |
| 7 | FS | 7 | DSH | Mixed tubular carcinoma | I | 21 | NA | No | |
| 8 | F | 5 | DSH | Tubulopapillary carcinoma | I | 12 | No | No | |
| 9 | FS | 10 | DSH | Intraductal papillary carcinoma | I | 17 | No | Yes | No |
| 10 | FS | 9 | DSH | Complex carcinoma | I | 17 | NA | Yes | No |
| 11 | FS | 5 | DLH | Tubular carcinoma | I | 12 | NA | No | |
| 12 | FS | 8 | DSH | Tubulopapillary carcinoma, cystic | II | 32 | No | No | |
| 13 | FS | 10 | DSH | Tubular carcinoma | II | 24 | NA | No | |
| 14 | FS | 10 | DSH | Tubulopapillary carcinoma | II | 10 | No | Yes | No |
| 15 | FS | 7 | DSH | Comedocarcinoma | II | 16 | No | Yes | No |
| 16 | FS | 7 | DSH | Tubulopapillary carcinoma | II | 40 | NA | No | |
| 17 | FS | 4 | DSH | Cribriform carcinoma | II | 15 | No | No | |
| 18 | FS | 13 | DSH | Simple adenocarcinoma | III | 4 | Yes | Yes | No |
| 19 | F | 8 | Persian | Tubulopapillary carcinoma | III | 8 | Yes | No | |
| 20 | FS | 11 | DSH | Simple carcinoma | III | 6 | Yes | Yes | Yes, to LN |
| 21 | FS | 10 | Siamese | Simple carcinoma | III | NA | Yes | Yes | Yes, to LN |
| 22 | FS | 10 | DSH | Simple carcinoma | III | 82 | Yes | No | |
| 23 | FS | 13 | DSH | Tubular carcinoma | III | 79 | No | No | |
| 24 | FS | 9 | DSH | Tubular carcinoma | III | 37 | NA | Yes | Yes, to LN |
| 25 | F | 6 | DSH | Tubular carcinoma | III | 9 | Yes | Yes | Yes, to LN |
| 26 | FS | 10 | Ocicat | Tubular carcinoma | III | 14 | Yes | Yes | Yes, to LN |
| 27 | FS | 11 | DSH | Tubular carcinoma | III | 68 | No | No | |
| 28 | FS | 9 | DSH | Simple carcinoma | III | 4 | Yes | No | |
| 29 | FS | 13 | DLH | Comedocarcinoma | III | 14 | Yes | No | |
| 30 | FS | 14 | | Solid and tubular carcinoma | III | 11 | Yes | Yes | Yes, to LN |
| 31 | FS | 12 | DLH | Tubular carcinoma | III | 26 | Yes | Yes | Yes, to LN |
| 32 | FS | 12 | | Cribriform carcinoma | III | 28 | No | No | |

NA: information not available
